# Human iEEG recordings during audiovisual speech violations support and extend predictive routing models

**DOI:** 10.64898/2026.10.30.760243

**Authors:** Jadyn V. Collado, Hamed Nejat, Vibhangini S. Wasade, William C. Stacey, David Brang, André M. Bastos

## Abstract

Rather than constantly reconstructing the world solely through sensory input, predictive processing theories posit that the brain utilizes predictions maintained as internal representations to guide sensory processing. An emerging model, predictive routing, proposes that an interplay between neuronal gamma (40-150 Hz) and alpha-beta (10-30 Hz) rhythms implements a dynamic mechanism for predictive processing. Namely, predictions enhance alpha-beta oscillations that suppress gamma activity during the processing of predicted inputs. While this model is supported by animal studies, its validity has not been fully explored within the human brain. The present study addresses this by leveraging human intracranial EEG (iEEG) recordings across superior temporal and prefrontal/premotor areas of seven presurgical epilepsy patients during an audiovisual speech paradigm. We calculated iEEG power in the theta, alpha-beta, and high gamma bands during simple auditory deviance, multisensory integration, and a condition where an incongruent visual cue (lip movements) clashed with an auditory stimulus (a syllable), creating audiovisual incongruence that can elicit the McGurk illusion. Results revealed increased high-gamma and decreased alpha-beta power in the superior temporal cortex in response to auditory deviance and the McGurk condition, while frontal areas, despite reflecting a similar decrease in alpha-beta power, did not present this high gamma prediction error signal. Instead, the theta band revealed a strong prediction error signal in the McGurk condition, consistent with theories on the role of theta in reorganizing cortical networks. These results both support and extend existing predictive routing models for human-specific forms of predictive processing.

**Significance Statement:** Predictive processing proposes that the brain relies on predictions which are created and maintained as internal representations of external reality, situating perception as an interaction between these expectations and new sensory input. Language provides the ultimate test of this model in humans, revealing how prediction shapes both basic sensory experience and high-level concepts. Although established work has supported distinct oscillatory signatures of predictive processing, much of the existing literature has relied on animal models or noninvasive human recordings. This study therefore aims to bridge this gap by leveraging intracranial EEG in human patients during audiovisual speech to investigate the underlying neural mechanisms of efficient prediction in the human brain.

## Introduction

Rather than perceiving the world solely through sensory input, predictive processing posits that the brain relies on predictions maintained as internal representations of expected reality (Bastos et al., 2012; K. Friston, 2008; Keller & Mrsic-Flogel, 2018; Rao & Ballard, 1999). When predictions differ from true sensory input, prediction error signals are generated to update expectations, reducing future error. Language provides a particularly revealing test of this model, demonstrating how prediction shapes sensory experience and high-level concepts (Arnal et al., 2011; Kutas & Hillyard, 1980).

Traditional predictive processing models propose distinct prediction and prediction error units (Bastos et al., 2012). Predictions in higher-order areas are propagated to lower-order areas and subtracted from ascending sensory input (Jiang et al., 2022) with the residual propagated up the cortical hierarchy as a prediction error to update representations (Chao et al., 2022). An alternative model, predictive routing, proposes that the brain calculates error more efficiently by proactively priming pathways associated with predicted stimuli, explaining prediction errors as the transmission of unexpected input through a pathway that was not primed by inhibition (Bastos et al., 2020; Gabhart et al., 2025; Xiong et al., 2024).

Established work has identified frequency-specific dynamics across these models, with alpha-beta oscillations linked to periods of prediction and high gamma activity associated with prediction error (Bastos et al., 2020; Chao et al., 2018; Dürschmid et al., 2016; Xiong et al., 2024). Theta oscillations have been further proposed to organize these error signals and coordinate top-down networks (Canolty et al., 2006; Han et al., 2025; Köster, 2024; Lakatos et al., 2005; Ruffini et al., 2025). Yet, characterizing these signatures demands the spatiotemporal resolution of intracranial recordings (Gabhart et al., 2025; Xiong et al., 2023). Much of the existing literature utilizes animal models (Chao et al., 2018) or noninvasive human work (Arnal et al., 2011; Arnal & Giraud, 2012; Chao et al., 2022), limiting insights into the fine-grained spatiotemporal dynamics of human cognition. This study therefore aims to investigate the predictive routing model through human intracranial EEG during an audiovisual speech paradigm, providing the ideal foundation to generalize mechanisms extensively studied in animal models to the human experience.

Within the auditory domain, high gamma responses are reliably enhanced in auditory regions during simple deviations from an expected sound and in frontal areas following violations (Dürschmid et al., 2016; Fuhrer et al., 2025; Nourski et al., 2018, 2021; Xiong et al., 2024). These frequency specific dynamics extend to multisensory audiovisual speech processing, where visual speech modulates high gamma auditory cortex activity prior to sound onset (Besle et al., 2008; Brang et al., 2020; Peelle & Sommers, 2015). At the spectral level, Karthik et al. (2021) further identified pre-auditory beta suppression and increased high gamma power in auditory areas during informative visual speech, followed by reduced theta power during later auditory processing relative to auditory only conditions.

The present study tests and extends these findings using a McGurk audiovisual speech paradigm, in which preparatory mouth movements (visemes) precede corresponding units of speech (phonemes) by approximately 150-200 ms (Arnal et al., 2011). By manipulating whether the preparatory visual cue matches or misinforms the later auditory signal, this paradigm allows for the separation of temporal windows of prediction and subsequent error in invasive human recordings. To investigate hypotheses of predictive routing, this study evaluates responses within prefrontal/premotor cortex and the superior temporal cortex (STG/STS), representing higher and lower-order regions implicated in generating predictions (Arnal et al., 2011; Fuster & Bressler,

2015) and audiovisual integration (Beauchamp et al., 2004), while also allowing for the isolation of visually driven predictive responses within an auditory processing region. Analyses focused on theta (2-6 Hz) alpha-beta (8-30Hz) and high gamma (70-150 Hz) activity. Accordingly, we hypothesized increased high-gamma activity during prediction error (H1) and condition-dependent modulation of pre- and post-stimulus alpha-beta (H2). Additionally, theta activity in the time-frequency representation motivated exploratory analysis of the theta band across conditions.

## Materials and Methods

Participants of this study were seven patients undergoing clinical epilepsy evaluation with intracranial EEG monitoring at the University of Michigan and Henry Ford Hospitals. The location of implanted electrodes was decided by patients’ clinical teams with no input from researchers, ensuring that our study posed no added risk to participants. Ethics and Institutional Review Board approval was obtained by the University of Michigan (HUM00118869) and Henry Ford Hospital (IRB 10955). Data collection from patients was anonymized for analysis, with four female and three male patients with an average age of 30.1.

## Experimental Design and Statistical Analyses

### Description of Task

To investigate the oscillatory dynamics of predictive processing in humans, this study analyzes intracranial EEG (iEEG) data collected during the McGurk paradigm, which leverages the natural lag between the preparatory facial movement to produce speech and the onset of sound itself to effectively separate the temporal window of prediction from the onset of prediction error (Arnal et al., 2011). Stimuli consisted of 2-second audiovisual clips in which a recording of a person mouthing a syllable (viseme) is played with an audio overlay (phoneme) of the same or a different syllable. In these audiovisual clips, the initial preparatory movement of the face began prior to the presentation of the auditory cue, allowing participants to form early predictions on which later auditory stimulus would be presented.

Audiovisual blocks were defined as the group of trials where a viseme and phoneme were concurrently presented, while auditory only blocks were those in which a phoneme was presented with no visual stimulus. In both audiovisual and auditory only blocks, the ‘standard’ phoneme was the sound which constituted the majority of trials, while the ‘deviant’ phoneme was the sound which was presented rarely. Counterbalanced blocks maintained the identical structure but switched the original standard and deviant phonemes. Thus, across the 192 total trials, 96 used ‘Ba’ as the standard phoneme and 96 used “Ga” as the standard phoneme.

Within each 96-trial block, 48 trials were auditory only and 48 were audiovisual. In auditory only blocks, the standard phoneme was presented in 75% of trials (72 trials total), while the deviant phoneme was presented in 25% of trials (24 trials total). In the Audiovisual blocks, for present analyses, audiovisual standard and McGurk trial results were analyzed. In audiovisual standard trials, a matching, standard viseme-phoneme pair is presented in 50% of trials. In McGurk conditions, a ‘deviant’ viseme is presented before the standard phoneme, functionally misinforming the later standard sound in 25% of audio-visual trials. Each auditory stimulus lasts 1000 ms and when accompanied by a visual stimulus, the visual cue begins 500 ms before sound onset after a 500 ms interstimulus interval. For all analyses, ‘pre-stimulus’ refers to the time period before the presentation of an auditory stimulus. This early visual onset provides a temporal window in which participants can form predictions based on viseme cues prior to the auditory input. The total experiment duration was therefore approximately 7 minutes given breaks between blocks (Figure 1).

**Figure 1:**
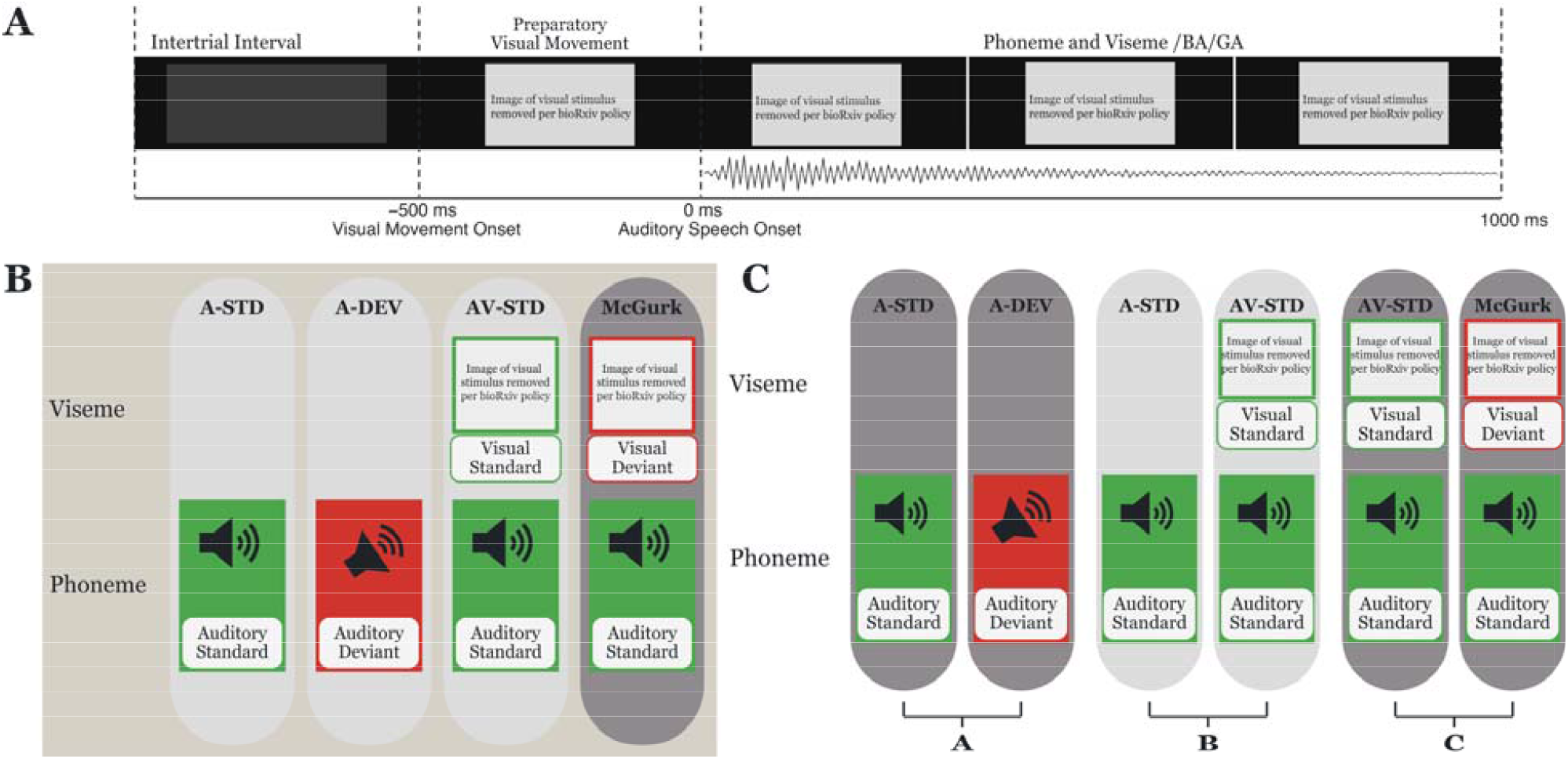
Experimental Paradigm. Each panel represents an element of the experimental paradigm including timing, trial types, and experimental contrasts. Panel A displays trial timing, consisting of a 2 second trial window beginning with a 500 ms intertrial interval period followed by the viseme presentation preceding auditory onset. Panel B displays the four analyzed trial types, with each rectangle representing a different condition type, determined by the pictured auditory and visual stimuli. From left to right, the trial types are auditory standard, auditory deviant, audiovisual standard, and McGurk. The top row of each rectangle represents the visual stimulus presented within this trial type, with the absence of a photo denoting there was no visual stimulus presented. The bottom row image of each rectangle represents the auditory stimulus presented within each trial type. In both visual and auditory stimuli depictions, the green squares indicate a standard stimulus and the red squares with a slanted speaker icon indicate a deviant stimulus. Panel C represents the experimental contrasts analyzed in the present study. From left to right, (A) Auditory Standard vs Auditory Deviant, (B) Auditory Standard vs Audiovisual Standard, and (C) Audiovisual Standard vs McGurk.

### Description of Behavioral Responses and Task Performance

Because data were collected over a 2.5-year period, the response type and instructions were refined across patients while the critical stimulus structure remained constant. Four patients performed a deviance-detection version, pressing a response button whenever they heard the deviant syllable and withholding responses to standards. Two patients performed a two-alternative forced-choice version, reporting the syllable they heard (/ba/ vs /ga/) with a button press after every trial within a 750-ms response window. One patient performed the same overt identification task, naming the syllable aloud after each trial. Forced-choice data from the two patients confirmed that the syllables and deviants were readily discriminable: when a response was registered within the response window, auditory identification was at or near ceiling, with greatest sound identification reported on McGurk trials. Button presses were not usable for the detection patients: a response-device logging error prevented registration of the response pad in the four patients tested. Because all analyses concern stimulus-locked neural responses and auditory deviance effects do not depend on an overt response, patients were included irrespective of the completeness of their behavioral records; the available identification data verify that patients perceived and could accurately categorize the stimuli.

### Electrode Placement

Prior to implantation, MRI images of each patient were taken with no additional identifying information to aid in electrode placement based on each patient’s clinical need by the University of Michigan and Henry Ford surgical teams. All experimental analyses utilize predetermined electrode placements which fell into areas of interest for the study. No additional electrodes were implanted for the purposes of this study. In a typical patient, approximately 3-7 multi contact electrodes were implanted with 4-16 contacts each, resulting in broad spatial coverage across cortical and subcortical areas. Electrode probes had 5 mm center to center spacing for University of Michigan patients and 3.5 mm spacing for Henry Ford patients. Preoperative T1 MRI and postoperative CT images were coregistered to the preoperative T1-weighted MRI using normalized mutual information in SPM (Friston, 2007); transformations were estimated without reslicing. Cortical surfaces were reconstructed with FreeSurfer. Electrode contacts were identified from CT intensity using custom software implementing the geometry-based method of Brang et al. (2016). For group analyses, each participant’s FreeSurfer reconstruction was affinely registered to the cvs_avg35_inMNI152 template using mri_robust_register, and the resulting RAS-to-RAS transformation was applied to each contact. STG/STS and prefrontal/premotor electrodes utilized for analysis were determined through this automated grouping followed by manual review of electrodes, ensuring patient-level differences in anatomy were accounted for. The electrode locations used in this study are visualized in Figure 2 on a template brain (cvs_avg35_inMNI152 template).

**Figure 2:**
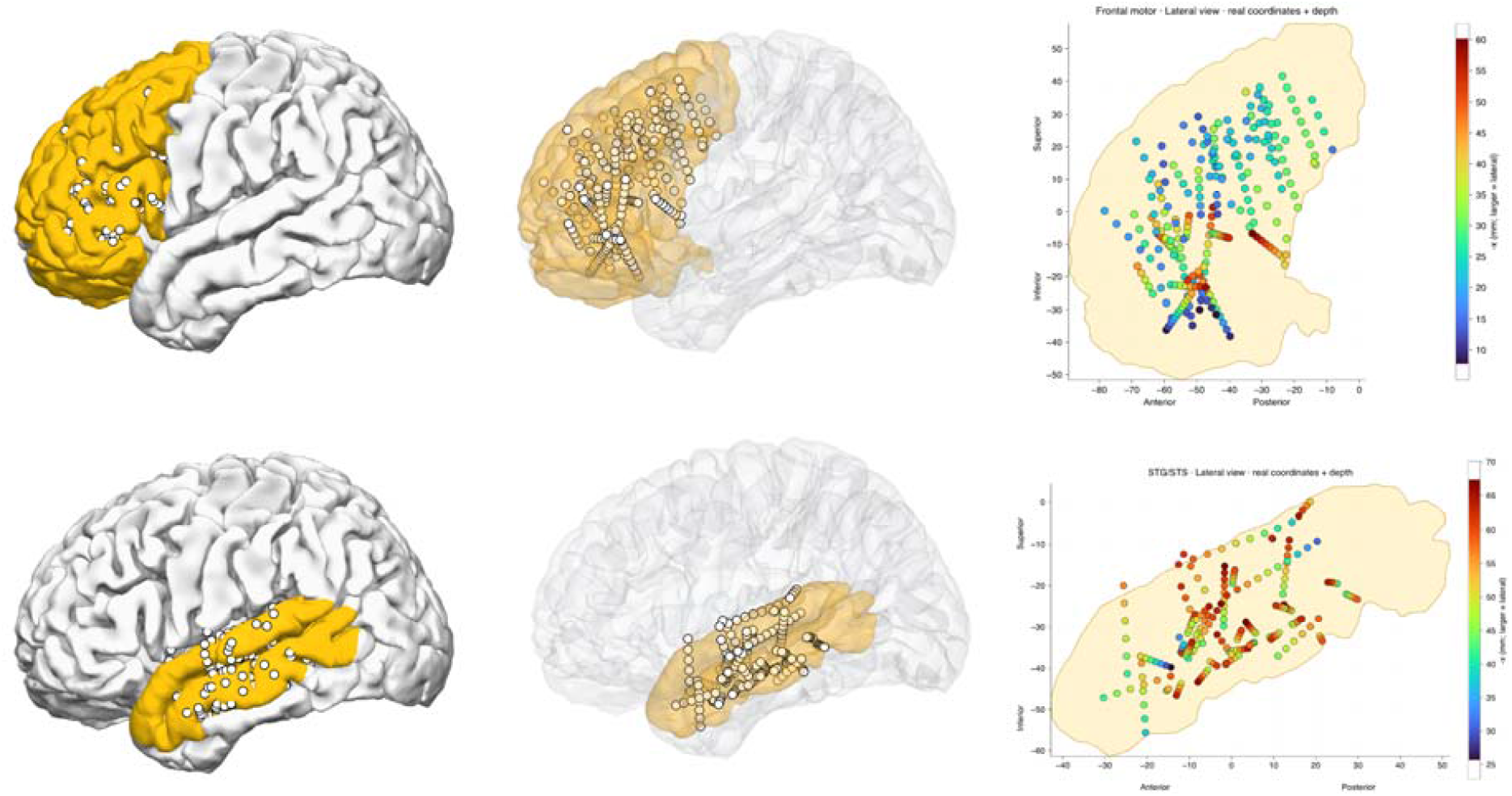
Electrode Placement and Regions of Interest. Each panel represents the approximate location of 470 electrode contacts across the two highlighted regions of interest across all patients projected into one common sample space. The top row displays all prefrontal and premotor electrodes while the bottom row displays all superior temporal gyrus and sulcus electrodes used in the following analyses. Across both rows, the leftmost column depicts a surface image of these contacts on an opaque brain template while the center column depicts all electrode contacts on a translucent brain template for visibility. The rightmost column depicts an orthographic projection of all electrode contacts within each region of interest colored by depth, with warm tones indicating more lateral and cool tones indicating more medial electrode placement.

## Data Analysis

All analyses were conducted using a spectral power analysis pipeline adapted from Brang et al. (2022), which has been applied in intracranial EEG studies of sensory processing. To reduce noise, Common Average Referencing (CAR) was utilized, in which each electrode channel is referenced to the average signal across all electrodes, displaying a voltage difference in reference to the mean (Huang et al., 2024). Filtering was then performed in MATLAB using custom scripts. A high-pass filter at 0.01 Hz was applied to remove extremely slow voltage drifts unrelated to neural activity or too slow to be the oscillations of interest (theta-gamma). A notch filter at 60 Hz and higher-frequency multiples was then used to eliminate electrical line noise from the recording environment (Brang et al., 2022). The sampling rate for this processing was determined at the hospital recording site as either 1024 Hz (n = 4) or 1000 Hz (n = 3), depending on hospital location, allowing for 2049 or 2001 sampling windows within each 2-second window per electrode for analysis. Experimental analyses of channels were limited to electrodes placed in cortical tissue in the prefrontal cortex (PFC, n = 247, see Table 1), including premotor areas, and superior temporal gyrus and sulcus (STG/STS, n = 223, see Table 1) regions. For each patient, neural responses were recorded from 192 trials, resulting in 384 total seconds of continuous data per electrode channel. This continuous data was divided into epochs, each including 500 ms preceding visual stimulus onset, 500 ms of visual stimulus presentation, and 1000 ms of auditory stimulus presentation, for a total duration of 2 seconds per epoch. Time was defined relative to auditory stimulus onset (0 ms), with -1000 ms marking the beginning of the epoch and 1000 ms marking the end of the auditory stimulus. This structure is maintained regardless of whether a visual stimulus was presented, allowing both intertrial and pre-stimulus activity to be measured and compared across condition types.

**Table 1.** Electrode counts per patient by region

| **Patient** | **STS/STG** | **Prefrontal/Premotor** |
|---|---|---|
| 1412 | 30 | 60 |
| 1401 | 27 | 48 |
| 1403 | 38 | 19 |
| 1404 | 27 | 29 |
| 1331 | 42 | 10 |
| 1396 | 20 | 49 |
| 1417 | 39 | 32 |
| **Total** | **223** | **247** |

*Note. STS/STG = superior temporal sulcus/gyrus.*

Morlet wavelet convolution was used to decompose each trial’s signal into its frequency components over time, allowing analysis of frequency-specific power and its temporal dynamics (Brang et al., 2022). Trial-averaged theta, alpha, beta, and gamma power were analyzed across all patients, with power expressed relative to a baseline period from -800 to -600 ms. This window falls within the intertrial interval prior to any visual stimulus onset, ensuring it was free of task related neural activity. For theta and alpha-beta analyses, 3-cycle wavelets were utilized, resulting in 500-1500 ms of temporal smoothing for analysis in the theta (2-6 Hz) range, 100-375 ms in the alpha-beta (8-30 Hz) range. For high gamma (70-150 Hz) analyses, 20-cycle wavelets were used, resulting in 133-286 ms of temporal smoothing within this frequency range. For each patient, the mean time-frequency power during a pre-stimulus baseline window (-800 to -600 ms) was compared to its activity during a post-stimulus window (100 to 400 ms) to identify responsive electrodes. Within each analyzed frequency range and region, electrodes with a z-score greater than 2.3 in any one condition type were retained, ensuring later analysis of task-relevant activity. To allow for grouped analyses, all patient-level data were resampled to the most common sampling rate using linear interpolation before pooling. Thus, data acquired at 1024 Hz (n = 4 patients) remained unchanged, while data acquired at 1000 Hz were upsampled to match this rate.

## Statistics

To compare power changes across specific frequency bands as a function of task epoch, we utilized a nonparametric, cluster-based pipeline adapted from Maris & Oostenveld (2007). This pipeline was utilized to control for the multiple comparisons problem. By using this cluster-based permutation model, significance is identified across a cluster of time points within a frequency band and then tested over a null distribution of randomly shuffled cluster values, thus reducing the odds of false positive values.

For each of the 2000 permutations, within each anatomical area and frequency band pooled across all patients, a subset of electrodes from the two analyzed conditions were randomly selected, and the corresponding rows (corresponding to trial averaged electrode activity) were swapped between the two condition matrices. Each row was exchanged as a complete time series, preserving electrode and sampling window order while randomly reassigning the condition label of the electrode. Between these two shuffled matrices, a 2-tailed dependent samples t-test was performed at each sampling window, resulting in a one row, 1024 column vector of t-values. Clusters were then determined by summing the concurrent t-values exceeding the critical threshold (|t| > tcrit, uncorrected p < 0.05), where tcrit = tinv(0.975, df). The number of included electrodes varied by analyzed area and frequency range. The largest cluster per shuffle was then saved and plotted on a histogram of the largest random clusters from all 2000 permutations to form a null distribution.

The real, unshuffled data was then evaluated with the same pipeline, but without shuffling the values to get the t-values and clusters of the real data. These real clusters were then compared to the null distribution generated from the shuffled data to determine whether they exceed the 95th percentile threshold. As an additional criterion, any prestimulus clusters passing the second level test for the auditory only contrast were used to establish an additional cluster threshold. Clusters in all following contrasts were included only if their summed t-values exceeded both the permutation-based 95th percentile threshold and the summed t-values of any significant prestimulus auditory only cluster. By placing the actual found clusters over this random distribution, we then know whether this found cluster has a greater than random chance of appearing proving that it is not simply the result of multiple comparisons

### Code accessibility

All following analyses used custom MATLAB scripts and a pipeline made in collaboration between the University of Michigan and Vanderbilt University available upon request.

## Results

All following analyses consisted of intracranial EEG recordings pooled across seven patients with electrode contacts located within prefrontal/premotor cortex (n = 247) and superior temporal gyrus/sulcus (n = 223) (Table 1). Each participant was presented 2-second audiovisual clips in which a recording of a person mouthing a syllable (viseme) is played with an audio overlay (phoneme) of the same or a different syllable depending on trial type. Audiovisual blocks were trial groups where a viseme and phoneme pair was concurrently presented, while in auditory only blocks, a phoneme was presented with no visual stimulus (Figure 1A).

To establish and extend frequency-specific responses, we analyzed three experimental contrasts: (1) Auditory only Standard vs, Auditory only Deviant, (2) Auditory only Standard vs Audiovisual Standard, and (3) McGurk vs Audiovisual Standard (Figure 1C). To isolate for the effects of interest, cluster-based nonparametric randomization tests (Maris and Oostenveld, 2007) were applied between the ERSP responses of contrasted conditions within the theta, alpha-beta, and gamma frequency ranges. Significant clusters are reported if they exceeded an alpha threshold of p< 0.05, corrected for multiple comparisons (see Methods).

### Group-Level Time Frequency Analyses

Figure 3 shows the time-frequency representation of Event Related Spectral Power (ERSP) for auditory only (Figure 3A), audiovisual standard (Figure 3B), and McGurk (Figure 3C) trial types, averaged across 223 STG/STS electrodes from all patients from 1-150 Hz. In response to auditory stimuli across these conditions, there is a distinct increase in power in the high gamma band as well as a decrease in the alpha-beta band relative to the baseline (-800 to -600 ms) (Figure 3A-C). In the McGurk condition however, there is also a concurrent decrease in alpha-beta and increase in theta power observed immediately before and following auditory stimulus onset which is prolonged in duration (Figure 3C). This representation, as well as prior established predictive processing work outlining the functional roles of these oscillations, provides justification for subsequent analyses of theta, alpha-beta, and gamma band activity across contrasted conditions.

**Figure 3:**
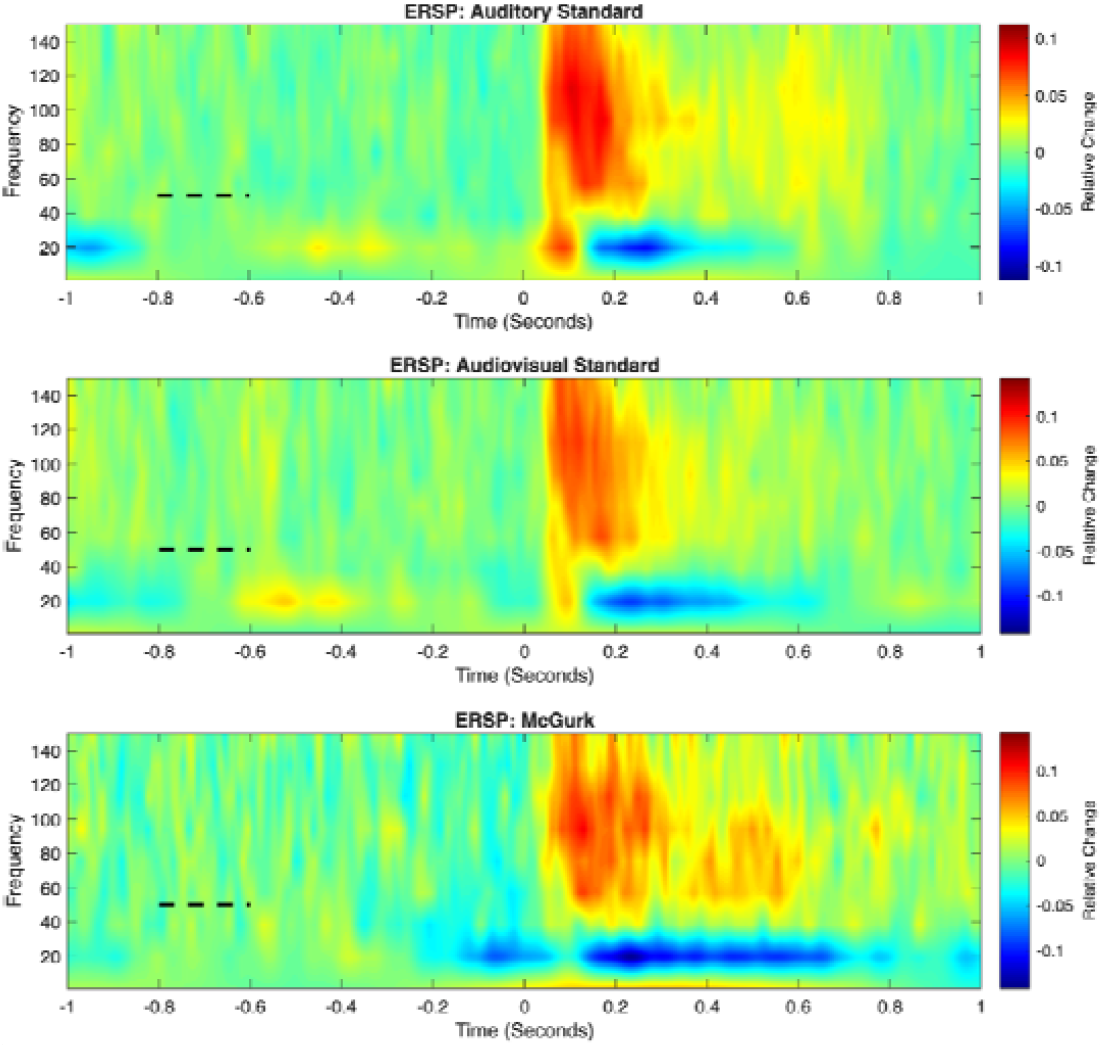
Distinct Theta, Alpha-beta, and Gamma Responses to Auditory Stimuli. Each panel shows the change in spectral power relative to the baseline period from -800 to -600 ms, averaged across all STG/STS electrodes and trials of the selected condition. Panel A displays auditory only standard trials, panel B audiovisual standard trials, and panel C the McGurk trials. Within each plot, the X axis represents time relative to auditory stimulus onset (0 ms), where the baseline period is denoted by the black dashed line. The Y axis represents frequency (Hz), and the heatmap values reflect relative change in spectral power from baseline, with warm tones indicating increased power and cool tones indicating decreased power.

### Simple Auditory Deviance Responses

To establish simple auditory response and sensitivity to auditory deviance without prior visual stimuli, we compared auditory only deviant and auditory only standard trial types across frequency bands of interest. Both auditory standard and deviant conditions presented a clear stimulus locked response in STG/STS areas, confirming robust auditory response through a distinct increase in power following auditory onset (0 ms) across frequencies (Figure 4A-C). Frontal responses were distinct from STG/STS dynamics, with responses across frequency bands both increasing and decreasing relative to baseline and relative to deviance vs. standard conditions (Figure 4D-4F).

**Figure 4:**
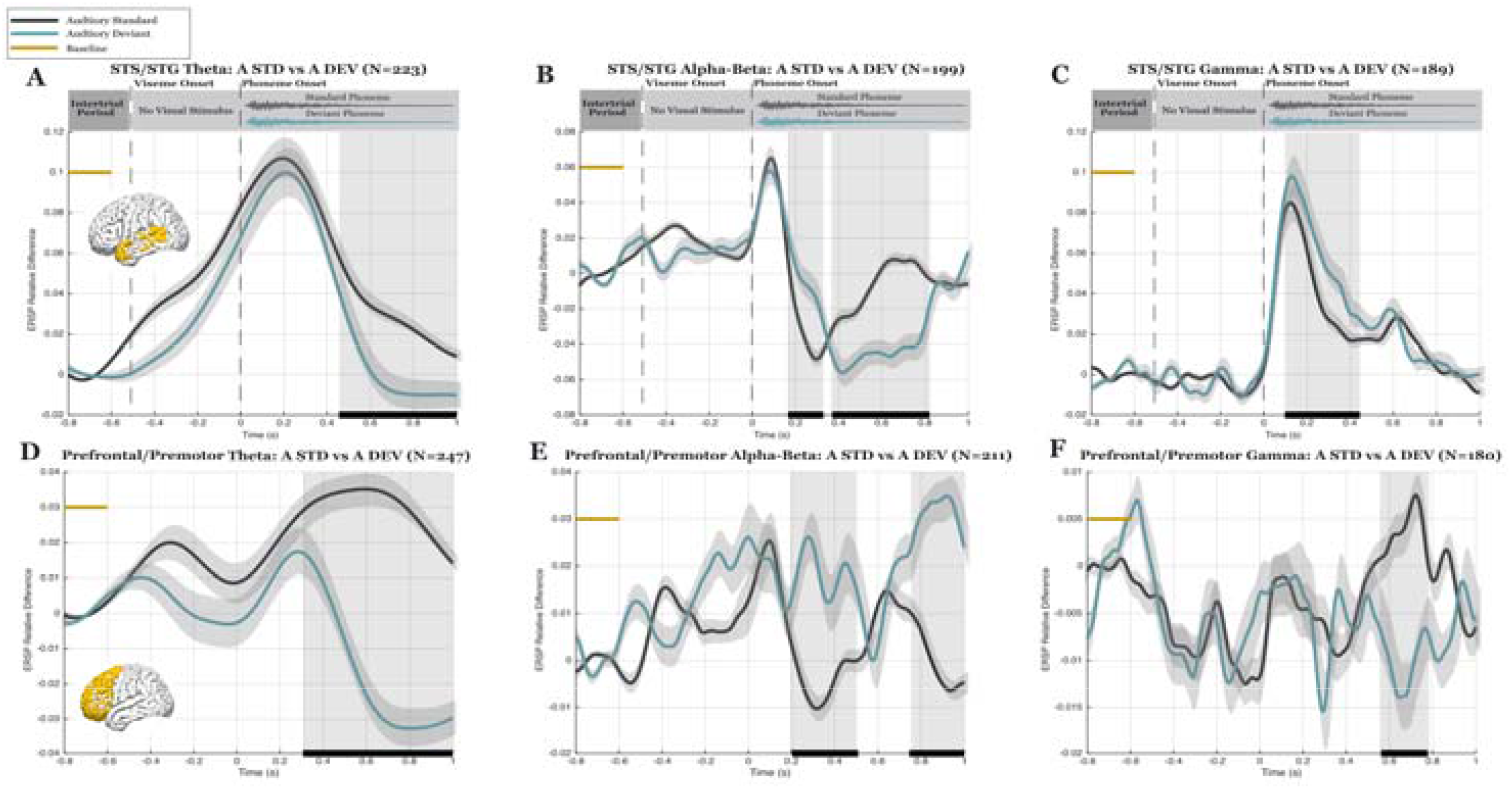
Relative Power Differences in Simple Auditory Deviance. Each panel shows trial and electrode averaged spectral power for auditory standard (grey plotted line) and auditory deviant (blue plotted line) trials. The top row (A-C) shows responses from STG/STS electrodes, and the bottom row (D-F) shows responses from prefrontal/premotor electrodes. Columns represent analyzed frequency bands (theta, alpha-beta, and gamma power from lef to right). The x-axis represents time relative to auditory stimulus onset (0 s), and the y-axis represents power relative to baseline (-800 to -600 ms). The baseline interval is indicated by the yellow horizontal bar. Black horizontal bars at the bottom of the panels accompanied by above grey shading indicate significant temporal clusters which exceed both the cluster forming t-threshold and the permutation based null distribution.

In the post-stimulus period (0 to 1000 ms), we examined whether there were significant differences in spectral power between the standard and deviant auditory stimuli to establish a response to simple auditory deviance. In STG/STS areas, the auditory deviant condition elicited significantly greater High Gamma Power (HGP) than the auditory standard condition from 0.097 s to 0.444 s (Figure 4C; (347.7 ms duration), Summed t-statistic (tcluster) = -1323.23, p < 0.001). Standard conditions initially elicited significantly lower power, before later deviant conditions elicited a later reduction in power relative to the standard condition type in the alpha-beta (Figure 4B; Cluster 1: time window: 0.165 s to 0.329 s (164.1 ms duration), Summed t-statistic (tcluster) = -690.77, p = 0.0185, Cluster 2: time window: 0.369 s to 0.820 s (451.2 ms duration), Summed t-statistic (tcluster) = +2565.09, p < 0.001 ) and theta (Figure 4A; time window: 0.455 s to 1.000 s (544.9 ms duration), Summed t-statistic (tcluster) = +1836.53, p = 0.0105) frequency ranges, reflecting distinct responses across frequency bands with HGP uniquely signaling simple deviance as supported in previously established work (Dürschmid et al., 2016; Fuhrer et al., 2025; Nourski et al., 2018, 2021; Xiong et al., 2024).

### Distinct Prefrontal Auditory Deviance Response

In prefrontal/premotor areas, as supported in prior literature (Dürschmid et al., 2016; Fuhrer et al., 2025; Nourski et al., 2021), the same distinct HGP response did not emerge following simple deviance. Rather, the auditory standard trial type evoked a significantly greater response in the theta (Figure 4D; time window: 0.307 s to 1.000 s (693.4 ms duration), Summed t-statistic (tcluster) = +5189.39, p < 0.001) and gamma bands (Figure 4F; time window: 0.562 s to 0.777 s (215.8 ms duration), Summed t-statistic (tcluster) = +845.47, p = 0.0050), while there were two points of significance in the alpha-beta band in which the auditory deviant condition evoked a significantly greater power (Figure 4E; Cluster 1: time window: 0.202 s to 0.509 s (306.6 ms duration), Summed t-statistic (tcluster) = -1179.40, p = 0.0055, Cluster 2: time window: 0.745 s to 1.000 s (254.9 ms duration), Summed t-statistic (tcluster) = -1583.95, p = 0.0015).

### Increased Alpha-beta and Gamma Power During Prediction

Now having established simple auditory deviance response in the gamma band, we next analyzed multisensory effects by comparing auditory only vs audiovisual trial types to evaluate the relative difference in response to auditory stimuli given the presence of a preceding visual cue.

Additionally, by focusing on the prestimulus period during which a preparatory visual cue was presented, this analysis tested whether spectral power changed in the auditory cortex during the time period in which preceding visual information could predict the later phoneme.

In STG/STS areas, there was a significant increase in relative alpha-beta power for the audiovisual conditions within the prestimulus period (Figure 5B; Time window: -0.685 s to - 0.444 s (240.2 ms duration), Summed t-statistic (tcluster) = -1240.36, p = 0.0015), consistent with the role of these alpha-beta band oscillations in representing or conveying sensory predictions. Additionally, this same prestimulus audiovisual significance was revealed in two distinct clusters in the gamma range (Figure 5C; Cluster 1: time window: -0.684 s to -0.522 s (161.1 ms duration), Summed t-statistic (tcluster) = -538.67, p = 0.0250, Cluster 2: time window: -0.194 s to -0.047 s (147.5 ms duration), Summed t-statistic (tcluster) = -578.86, p = 0.0180), which reproduced prior work on multisensory integration (Brang et al., 2020). The inverse pattern was presented but nonsignificant in the theta range: greater theta power during auditory only conditions within the prestimulus period, while frontal areas showed significant prestimulus differences in relative power between contrasts in the theta band. Theta prefrontal dynamics displayed relatively less power in the audiovisual condition (Figure 5D; -0.672 s to - 0.128 s (543.9 ms duration), Summed t-statistic (tcluster) = +2287.78, p = 0.0040), mirroring nonsignificant lower level dynamics within this frequency range. Together these findings provide support for hypothesis two in auditory areas: increased alpha-beta power during the presentation of predictive information about the later phoneme, consistent with the role of these oscillations in representing or conveying sensory predictions. Additionally, these dynamics were supported in the gamma range within auditory areas, presenting increased HGP during the presentation of a visual stimulus.

**Figure 5:**
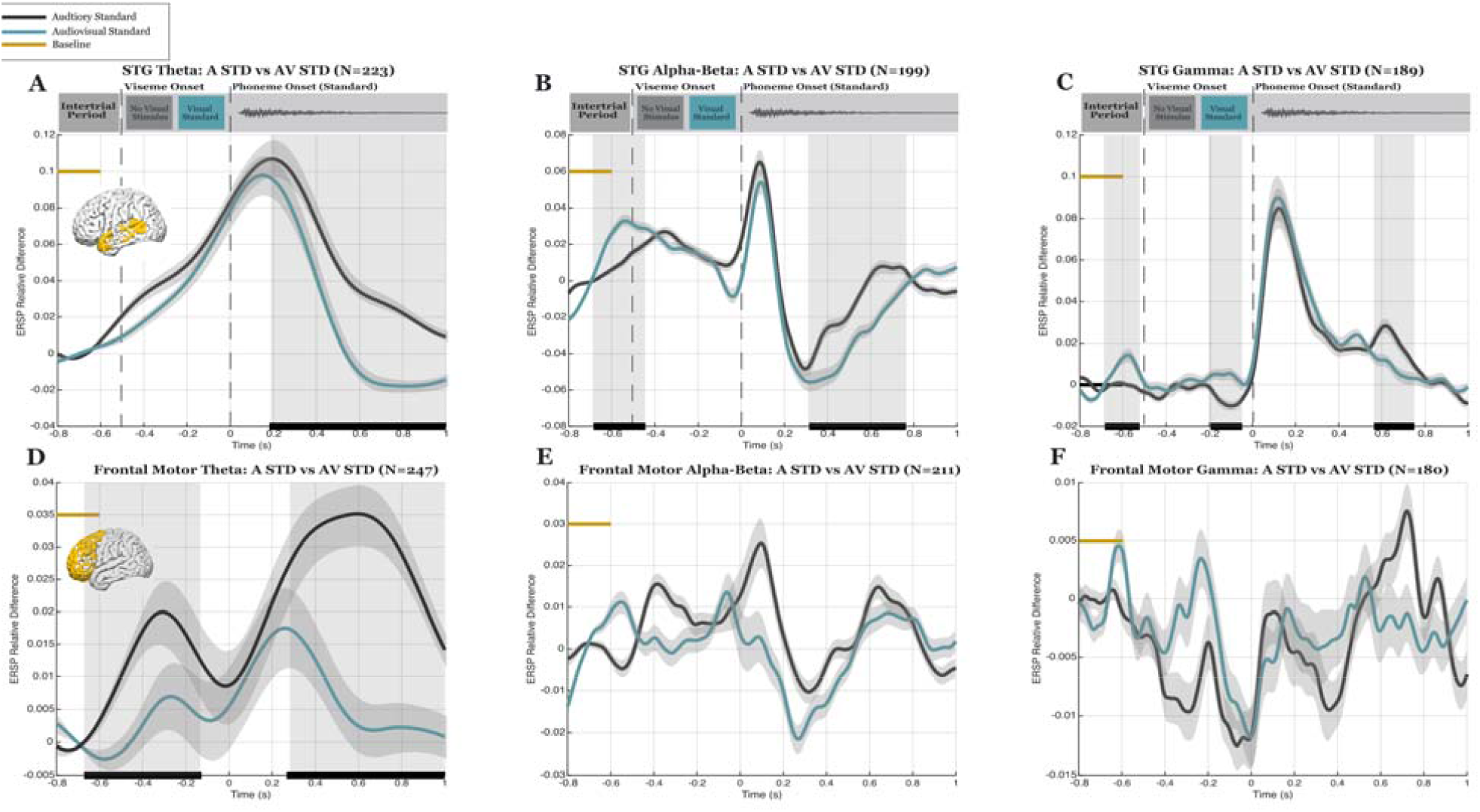
Relative power differences in multisensory deviance. Plot organization and conventions are as in Figure 4. Lines show trial and electrode averages spectral power for auditory only (grey plotted line) and audiovisual standard (blue plotted line) conditions across prefrontal/premotor (D-F) and STG/STS (A-C) electrodes. Black horizontal bars below grey shading indicates significant temporal clusters between conditions.

### Modulation of Auditory Processing Based on Informative Visual Cue

In the post stimulus period, informative visual information modulated post stimulus dynamics in the auditory cortex across theta, alpha-beta, and high gamma analyses. Across these frequencies, there was a significantly greater power in the post stimulus period to the auditory only condition in which no prior visual stimulus was presented relative to the audiovisual condition. This difference emerges first and was longest sustained in the theta band (Figure 5A; time window: 0.182 s to 1.000 s (818.4 ms duration), Summed t-statistic (tcluster) = +6319.05, p< 0.001), followed by the alpha-beta band (Figure 5B; time window: 0.314 s to 0.764 s (449.2 ms duration), Summed t-statistic (tcluster) = +2479.14, p< 0.001), before emerging last and the most briefly in the gamma band (Figure 5C; time window: 0.563 s to 0.751 s (187.5 ms duration), Summed t-statistic (tcluster) = +744.17, p = 0.0085). In frontal areas, there was only a significant relative increase in power for the auditory only condition in the post stimulus period in the theta band (Figure 5D; time window: 0.267 s to 1.000 s (733.4 ms duration), Summed t-statistic (tcluster) = +4771.59, p < 0.001), reflecting a relative increase in power during the presentation of an auditory cue without an early predictor. These results reveal increased activity across frequency bands in auditory and higher-order areas during the presentation of an auditory cue with no prior predictive visual information. This is consistent with prior studies that have shown a reduced magnitude of neuronal response to audio-visual speed compared to auditory-only speech (Simon and Wallace, 2018).

### Spectral Responses to Visual Deviance in the McGurk Condition

We next analyzed the McGurk condition. In McGurk trials, a deviant visual stimulus is presented to misinform the prediction of the later standard auditory cue (Figure 1B). During the presentation of the viseme and before auditory onset, McGurk trials exhibited reduced alpha-beta power across both STG and frontal regions (Fig 6B; Cluster 1: time window: -0.668 s to -0.410 s (257.8 ms duration), Summed t-statistic (tcluster) = -1230.66, p = 0.0040, Cluster 2: time window: -0.361 s to 0.243 s (604.5 ms duration), Summed t-statistic (tcluster) = -2820.18, p = 0.0005, Figure 6E; time window: -0.192 s to 0.009 s (201.2 ms duration), Summed t-statistic (tcluster) = -918.33, p = 0.0110). This points to a common prestimulus increase in alpha-beta to the presentation of a standard (predictable) visual cue relative to the deviant McGurk visual stimulus.

**Figure 6:**
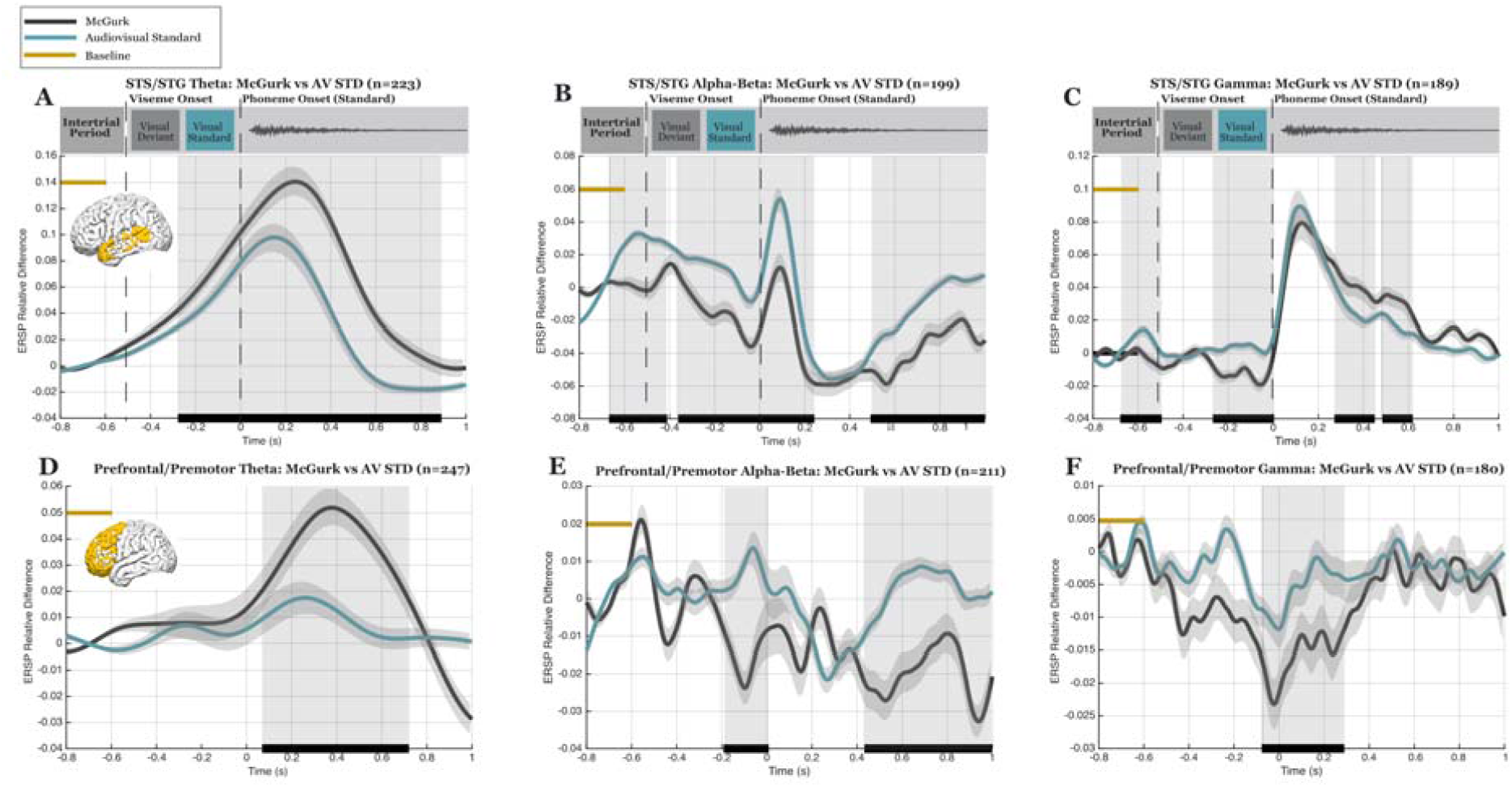
Relative power differences in the McGurk effect. Plot organization and conventions are as in Figure 4. Lines show trial and electrode averages spectral power for McGurk (grey plotted line) and audiovisual standard (blue plotted line) conditions across prefrontal/premotor (D-F) and STG/STS (A-C) electrodes. Black horizontal bars below grey shading indicates significant temporal clusters between conditions.

### McGurk Condition Evokes Prediction Error at Gamma Frequencies

We next examined how response to the same, standard auditory cue is modulated by prior visual information that either correctly or incorrectly informs the later auditory stimulus. Based on prior predictive processing frameworks, we hypothesized that mismatch between predicted and actual inputs would drive an increase in HGP following the presentation of an unexpected auditory stimulus in both auditory and higher-order regions.

In STG areas (Top row of Fig. 6), the McGurk condition evoked relatively increased gamma activity, emerging at two points in the post-stimulus period from 0.271 s to 0.619 s, (Figure 6C; Cluster 1: time window: 0.271 s to 0.450 s (179.7 ms duration), Summed t-statistic (tcluster) = +514.10, p = 0.0335; Cluster 2: time window: 0.483 s to 0.619 s (135.7 ms duration), Summed t-statistic (tcluster) = +413.62, p = 0.0495). However, the inverse direction of significance was found in frontal areas, with a significant decrease in frontal HGP from -0.077 s to 0.290 s (Figure 6F; time window: (367.2 ms duration), Summed t-statistic (tcluster) = -1022.24, p = 0.0035) These results signal region specific responses in which high gamma power is selectively increased during McGurk prediction error within auditory areas but decreased within the frontal cortex. These findings in the gamma band provide support for hypothesis one within auditory areas: increased high gamma power for prediction error or mismatch between predicted and actual inputs within the McGurk trial types, with the inverse response in frontal regions.

### Distinct Theta Responses During Audiovisual Incongruence

Although hypothesis one specified differences in high gamma dynamics during prediction error, the time-frequency representations in Figure 3C also provided insight into distinct activity in the Theta range during the McGurk condition. Given previous links between theta and gamma oscillations (Canolty et al., 2006; Köster, 2024; Lakatos et al., 2005; Ruffini et al., 2025), we also investigated whether these prediction related differences were present in the theta range. We found a widespread prediction error response in the McGurk condition relative to audiovisual standard trial types in the theta band (Figure 6A; 6D). There was significantly increased theta band power in the McGurk condition in both STG/STS and frontal areas (Figure 6A; STG/STS, time window: -0.278 s to 0.892 s (1169.9 ms duration), Summed t-statistic (tcluster) = +5405.19, p < 0.001, Figure 6D; frontal, time window: 0.068 s to 0.721 s (652.3 ms duration), Summed t-statistic (tcluster) = +2896.15, p = 0.0050). These results revealed signatures of a ‘pure’ prediction error-like response in frontal areas, in which the change in power relative to baseline only increased in the McGurk condition while audiovisual standard power remained relatively flat and close to baseline. Together these findings in the theta band provide additional support for hypothesis one in auditory and frontal areas. There was increased high gamma power for prediction error within the McGurk trial types in auditory areas.

## Discussion

### Summary of Findings

By leveraging human iEEG during the McGurk paradigm, this research enabled the separation of the temporal windows of prediction and error within STG/STS and prefrontal/premotor areas across seven human subjects. Initial analyses first established that patient recordings reproduced existing literature on simple auditory deviance and audiovisual integration. We observed a significant increase in high gamma power to simple auditory deviance in STG/STS areas without a corresponding significant frontal response. Predictive visual information also altered auditory region activity before sound onset and modulated later auditory responses across multiple frequency bands. When visual prediction was violated in McGurk trials, high gamma power increased in STG/STS without a significant corresponding increase in frontal cortex. However, the theta band revealed a ‘pure’ prediction error signal that spanned auditory and frontal areas.

These results, while consistent with key predictions of the predictive routing hypothesis, suggest that oscillatory signatures of predictive routing may be more context and region dependent than initially anticipated.

### Simple Auditory Deviance Response

Simple auditory deviance elicited a significant increase in high gamma power within the auditory cortex without a corresponding frontal high gamma response. These response profiles provide support for the distinct role of the gamma band in signaling simple prediction error, consistent with established findings that simple auditory deviance produces increased local high gamma power that may not reach higher levels of the cortical hierarchy (Dürschmid et al., 2016; Fuhrer et al., 2025; Nourski et al., 2021). Theta showed the opposite pattern, with a significant decrease in power for the auditory deviant relative to the standard trials across both auditory and frontal regions.

Alpha-beta responses were less uniform: following the presentation of the auditory stimulus, there was a brief early decrease in power for standard trials followed by a late decrease in power during deviant trials within STG, whereas frontal regions only showed greater power during deviant trials. These opposing patterns may reflect coordinated low and high frequency dynamics during simple auditory deviance, with a gradual decrease in alpha-beta power in STG/STS occurring alongside an increase in high gamma activity followed by a later reduction of theta power. Within the predictive routing framework, the later alpha-beta power reduction during deviant trials may represent diminished priming of the expected pathway, allowing unexpected sensory input to be expressed through increased high gamma activity within auditory areas.

However, frontal areas which did not show the same reduction in alpha-beta for deviance, also did not have a corresponding high gamma deviance response.

### Audiovisual Integration

Results supported existing audiovisual multisensory work, finding that the presentation of informative visual information modulated the later response to the same auditory stimulus (Besle et al., 2008, Simon & Wallace, 2018), while also replicating the frequency specific pattern reported by Karthik et al. (2021): greater theta and beta power during auditory only trials paired with a prestimulus enhancement of high gamma power in the audiovisual condition. We further extended this work with a longer poststimulus analysis window, revealing a later increase in gamma power during auditory only standard trials, as well as a prestimulus increase in alpha-beta power during audiovisual trials.

Supporting hypothesis two, there was a significant, early increase in alpha-beta and high gamma power in auditory areas during the presentation of informative visual information. Later following the auditory onset however, this relationship reversed: trials without the prior informative visual cue (auditory standard trial types), displayed a significant increase in power across areas and frequency bands. Together these findings suggest that visual prediction alters subsequent auditory processing, based on whether this stimulus was meaningfully predicted before its arrival.

### The McGurk Condition Reveals a Distinct Oscillatory State

With established simple deviance and multisensory responses, the McGurk condition provided a test of how these oscillatory dynamics change when an informative visual prediction is subsequently violated. Prior to auditory onset, where the informative visual stimulus was either standard (audiovisual standard) or deviant (McGurk), there is support for condition-specific anticipatory patterns across frequency bands: during the presentation of the visual stimuli, there was a significant decrease in power for the deviant McGurk visual stimulus relative to standard in both the alpha-beta and gamma bands across STG/STS and frontal areas. This gamma result then inverts following the auditory cue, with a significant increase in high gamma power for the McGurk conditions within STG areas. Because McGurk and audiovisual standard trials contained the same auditory stimulus, this increase in post stimulus power for the McGurk trial reflects the influence of a misinformative prior visual prediction on later auditory processing.

The timing of these effects is consistent with Arnal et al. (2011), who reported increased high gamma power associated with audiovisual prediction invalidation from approximately 350-500 ms following auditory onset, aligning with the effects observed in the present analysis. This significant response functions to support hypothesis one, high gamma power for prediction error in auditory areas locally, but this effect was not observed in frontal areas.

Instead, analyses of the theta frequency band revealed the broader prediction error response initially hypothesized in the gamma band, with a relative increase in theta power for the McGurk conditions spanning auditory and frontal areas. In the auditory only standard vs deviant as well as the audiovisual standard comparisons, theta consistently displayed a relative increase in power for the auditory only standard condition type, signaling that this band did not simply mirror high gamma dynamics. These results across contrasts thus signal that this increased theta power in the McGurk condition may not be the byproduct of a generic sensory response or a uniform reflection of gamma activity, but potentially a unique response to a higher level multisensory prediction error that is reflected in local auditory and higher-order prefrontal, premotor areas.

### Theta in Predictive Routing

Prediction error results in the theta band prove consistent with rising theories that the theta band may function to organize or carry faster local activity across cortical regions (Canolty et al., 2006; Köster, 2024; Lakatos et al., 2005; Ruffini et al., 2025). In the present study, localized high gamma prediction error response in auditory areas paired with a broader theta response spanning auditory and frontal areas raises the possibility that theta contributes to the integration of prediction error across the cortical hierarchy. These findings raise the question of whether the theta band is integrated into the larger predictive processing framework, acting as a functional vehicle of the prediction errors initiated in the gamma band during higher-order prediction errors.

### Limitations

Several limitations constrain the interpretation of these findings. First, recordings were obtained across a relatively small sample of patients with electrode placement determined solely by clinical need. As a result, the electrode location and count varied across participants, while the sample size limited reliable mixed effects modeling and led all analyses to rely on pooled data. Second, the present analysis characterizes frequency specific power changes rather than directional communication, laminar dynamics, or cross-frequency coupling. This work prompts future analyses to directly examine the interaction of these low and high frequency dynamics, while larger intracranial samples would permit mixed effects modeling to better account for individual variability.

### Summary & Predictive Routing Support

The present findings are consistent with predictive routing hypotheses: that predictions alter oscillatory state, particularly in the alpha-beta band, before sensory input arrives and that violations of prediction are expressed through distinct high gamma responses. Across both simple auditory and audiovisual conditions, prediction error was associated with increased high gamma activity in STG/STS, supporting a local sensory error signal within auditory areas.

Conversely, predictive visual information altered prestimulus activity before auditory onset and modulated later response to the same auditory stimuli, consistent with the idea that prior predictions proactively inform processing of future information. At the same time, our results suggest that the low and high frequency prediction and error signaling may be more flexible than originally proposed, considering the integration of the theta band into the predictive routing model in instances of higher-level prediction error.

In both simple deviance and McGurk analyses, we observed a push-pull relationship between alpha-beta and gamma power: a decrease in alpha-beta power in one condition was also reflected as an increase in high gamma power for the same deviant condition, potentially reflecting a distinct interplay between these oscillations in the production of local error. The theta band, however, consistently displayed increased power to the auditory only standard condition (Figure 4 and 5A), not signaling the same dynamic shifts as the alpha-beta and gamma bands until the McGurk condition was presented (Figure 6A). This pattern may suggest that during more complex, higher level errors, theta activity may contribute to coordinating prediction-related responses across regions, potentially understanding the multisensory integration aspects of the McGurk effect as shifting the frequency signature of error from the high-gamma to the theta band.

## Acknowledgements

This research was supported by the National Science Foundation (NSF) Faculty Early Career Development Program (CAREER) grant 2339210 (AMB) and the National Institutes of Health (NIH) NIDCD R01DC020717 (DB), and NIH R01NS094399 (WCS). We thank Ahmed Nawaz for assistance with data analysis.

The authors declare no competing financial interests

